# Fruit bromelain derived peptides destabilize growth of amyloidal fibrils

**DOI:** 10.1101/2020.04.20.051193

**Authors:** Sromona Das, Sangita Dutta, Ramesh K Paidi, Subhas C Biswas, Umesh C Halder, Debasish Bhattacharyya

## Abstract

β-Amyloid deposition as fibrillar plaques in brain is the primary cause of Alzheimer’s disease. We report potency of cysteine protease ‘fruit bromelain’ from pineapple in destabilising Aβ fibrils. Bromelain peptide pool (Mw<500 Da) obtained mimicking human alimentary tract digestion inhibited fibrillation from monomeric and oligomeric states of Aβ and irreversibly dissociated preformed fibrils into small oligomers of varied sizes. Time kinetics was followed by Thioflavin-T assay and microscopic imaging. Synthetic bromelain peptides corresponding to Aβ sticky region found using ClustalW analysis revealed specificity of peptides in destabilisation of amyloidal structures. Spectra of different molecular states of Aβ obtained from application of 8-anilino-1-naphthalenesulfonic acid, circular dichroism and Fourier-Transformed Infrared spectroscopy collectively indicated interaction dependent structural change. Probable mechanism for fibril dissociation was thus predicted. Peptides relieved Aβ cytotoxicity on pheochromcytoma cells and dissociated plaques in AD-type rats prepared by bilateral intracerebroventricular administration of Aβ in rat brain cortex. Pineapple being a phytoceutical, its efficiency to disaggregate amyloid bodies warrant further investigation.

**GRAPHICAL ABSTRACT:** 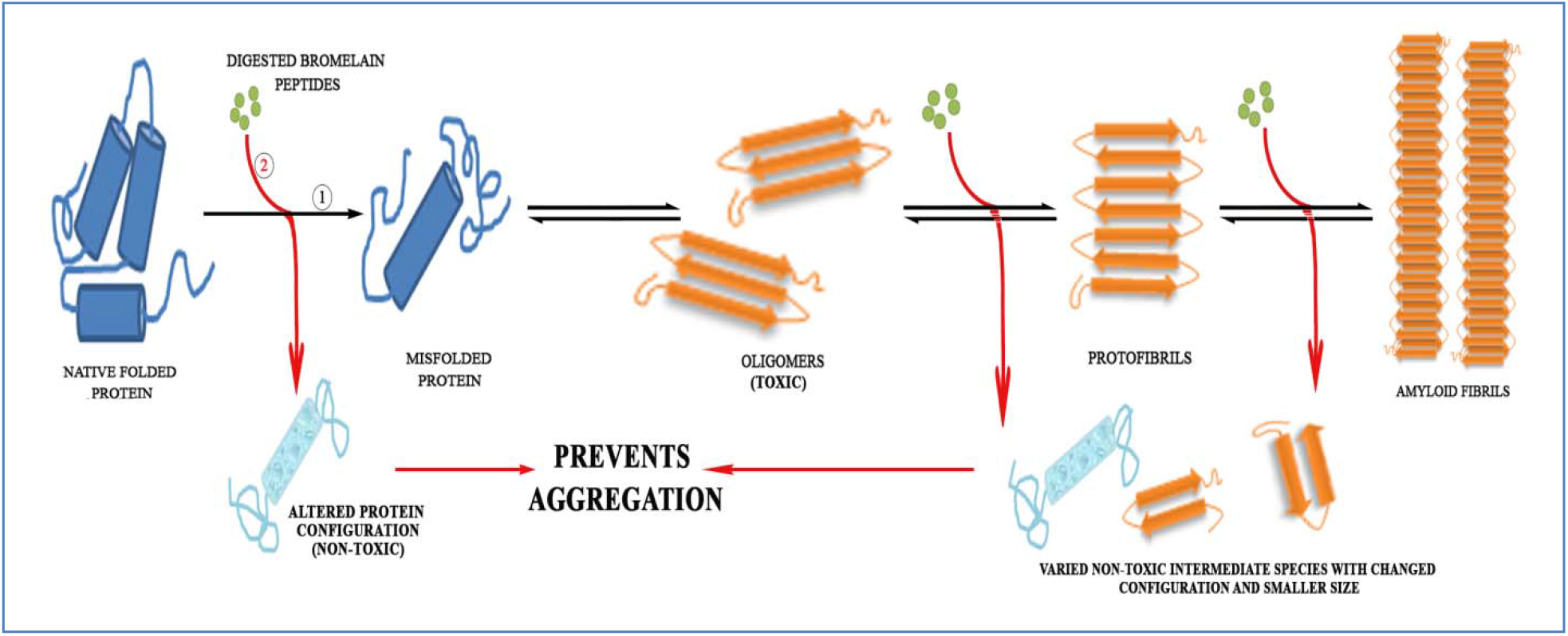

## INTRODUCTION

The occurrence of dementia has increased to one individual every 3 seconds, with 50 million affected people in 2018 and according to World Alzheimer’s Report, this will almost double every 20 years, reaching 82 million in 2030 and 131.5 million in 2050. Alzheimer’s disease (AD), the most widely studied and common form of dementia, accounts to 60–80% cases and is the sixth leading cause of death in USA **[1]**. Casual factors for AD though known for a very small 1-2% of the total population, it is still unknown for majority of cases. Though several factors, such as the ApoE4 genotype and polymorphism in several genetic loci have been identified alongside type2 diabetes, brain injury, stroke, diet and other environmental factors, aging stands as the first risk factor.

AD is an age-related progressive, neurodegenerative disorder characterized by gradual memory loss, cognitive abilities, characteristic neuropathological amyloid plaque depositions, formation of neurofibrillary tangles and severe neuronal loss in diverse regions of brain cortex **[2]**. A characteristic hallmark includes irreversible brain degeneration in elderly people. The amyloid deposits and senile plaques mainly contain insoluble, aggregated proteins, the main constituent being β-amyloid (Aβ42), derived from a 170 kDa cellular amyloid precursor protein (APP) **[3, 4]**. Several lines of evidence suggest a pathogenic role for Aβ assembly in progression of AD, and increasing evidences point specifically towards soluble protofibrillar intermediates as a pathogenic species **[5]**. Thus, there is considerable interest in studying the structures and assembly mechanisms of proteins into amyloid structures and their precursors. Consequently, though there is more than one stage that can be targeted, preventing aggregation is the primary therapeutic target.

Several pioneering work has led to the possibility of generating small and soluble oligomeric forms of Aβ *in vitro* **[6]** and these oligomers have proved detrimental on binding to synaptic neurons **[7]**. AD, a multifaceted disease involves multiple cellular changes, including synaptic and neuronal loss, activation of microglia and astrocytes, mitochondrial damage, and tau phosphorylation. Thus, various therapeutic strategies have been developed based on these modifications but no certain cure is available till date and is also nevertheless dose-dependent. Further, there are no available biomarkers that can relate symptoms of AD in individuals.

Physical exercise and healthy diets, few of the best remedies targeted towards good health decelerates growth of AD in elderly people, thereby aiding memory and learning skills in early AD patients and individuals having low cognitive impairment **[8]**. Natural products, a major component of healthy diets have multiple health benefits, including anti-inflammatory, antioxidant, anti-arthritis, neuroprotective and memory cognitive functions **[9]**. Many well known natural products and herbs currently include green tea, β carotene, vitamins E and C, rosemary, curcumin, ginseng, sage and many others **[8, 9]**. Concentrating specifically on natural products and targeting small molecules, we studied protective effects of pineapple extract derived enzyme, bromelain against Aβ induced toxicities in AD pathogenesis.

Bromelain, the cysteine protease from pineapple has broad specificity. It hydrolyzes diverse substrates like native and partly denatured collagen, elastin, casein, fibrin, hemoglobin etc. It offers a wide range of therapeutic efficacies and due to its efficiency after oral administration, safety and lack of undesired side effects; bromelain is being increasingly accepted as a phytotherapeutical drug **[10].** In this perspective, investigation was carried out to verify whether bromelain derived peptides could destabilize preformed Aβ aggregates. As a first step towards understanding interactions between fruit bromelain derived peptide pool and Aβ aggregate, we described effect of bromelain on preformed Aβ aggregates *in vitro*. Although structural information available for Aβ complexes is sparse, this analysis is likely to reveal features relevant to Aβ binding and generate initial hypotheses highlighting effect of peptide pool on Aβ aggregate. Earlier reports show that inactive and autodigested fruit bromelain act as a kinetic/nucleation inhibitor of amyloid formation in human insulin **[11].**

In this study, amyloid aggregates were reconstituted *in vitro* in a controllable manner and anti-amyloidogenic effect of bromelain peptide as an early quick battery test was performed before cellular and animal studies. The work described that fruit bromelain derived peptides, obtained from extensive digestion by various proteases found in human gastrointestinal tract, have an anti-aggregation property and can permeate the blood brain barrier (BBB) with respect to its small size. Further, three methods have been used to understand the effect in preventing formation of aggregates from (i) monomers, (ii) oligomers and also (iii) disaggregation of preformed Aβ fibrils. The self-assembling evaluation of Aβ *in vitro* will provide an opportunity to screen molecules for anti-amyloidogenic property of therapeutic significance for new AD drug discovery.

## EXPERIMENTAL SECTION

### Materials

Fine chemicals were procured as follows: Thioflavin T (Th T) from Acros Organics, Belgium; 8-anilino-1-naphthalene sulfonic acid (ANS), 1,1,1,3,3,3-Hexafluoroisopropanol (HFIP) and Paraformaldehyde (PFA) from Sigma–Aldrich, St. Louis, Missouri, USA; Dimethyl sulfoxide (DMSO) from HiMedia Laboratories, Mumbai, India; trypsin (3x crystallized, bovine pancreas), carboxypeptidase (bovine pancreas, type II – PMSF treated aqueous suspension), α-chymotrypsin (hog pancreas), elastase (porcine pancreas) and pepsin from SRL, Mumbai, India; acetic acid, formic acid, Acetonitrile (ACN) and Trifluoroacetic acid (TFA) from Merck, Germany; urethane from AMRESCO, Texas, USA; uranyl acetate from BDH Corporation, Mumbai, India; copper grids (300-mesh size) and mica sheets (size 20620 MM, 0.27 – 0.33 mm thickness) from Electron Microscopy Sciences, Pennsylvania, USA; MTT from Molecular Probes, Invitrogen, USA; Dulbecco’s Modified Eagle’s Medium (DMEM), penicillin-streptomycin, trypsin-EDTA and Fetal Bovine Serum (FBS) from Gibco, Maryland, USA. HPLC-purified peptides Aβ40/42 were procured from American Peptide, Sunnyvale, CA, USA. Physical and chemical homogeneity of monomeric Aβ peptide was verified by mass spectrometric analysis where other than trace amount of dimer, trimer and tetramer of the peptide, no other impurities could be detected.

### Preparation of Aβ40/42 oligomers and fibrils

Aβ40/42 oligomers and fibrils were prepared following Barghorn et al., 2005 **[12]**. Briefly, lyophilized Aβ peptide was reconstituted in 100% HFIP to a concentration of 1 mM. HFIP was removed by evaporation in a Speed Vac and then resuspended to 5 mM in anhydrous DMSO. This stock served as monomeric Aβ and was stored at −80°C. The stock was diluted to 400 μM with Phosphate Buffer Saline (PBS) containing 0.2% SDS and incubated at 37°C to form oligomeric intermediates. A further dilution with PBS to 100 μM and incubation at 37°C for 96 hr – 7 day formed amyloid structures having increased level of crosslinks.

### Preparation of fruit bromelain

Fresh ripe pineapple (100 g) was cut into small portions, smashed by a household grinder and centrifuged at 10,000 rpm for 15 min at 4°C to separate fibrous materials from edible portion. The lyophilized supernatant was reconstituted in 10 mM Na-phosphate, pH 7.5, and centrifuged as stated. The clear supernatant (2 ml) was passed through Sephadex G-50 column (1.5 × 90 cm) pre-equilibrated with the same buffer at 25°C. Flow rate was maintained at 12 ml/hr. Fractions collected were followed at 280 nm and assayed for proteolytic activity using azocasein as substrate **[13].** Proteolytic activity of bromelain was confined to the first peak fraction while the second contained pigments, salts and small peptides. Active fractions were pooled and dialyzed against 1 mM Na-phosphate, pH 7.5 and lyophilized. All spectrophotometric measurements were taken in Specord 200 spectrophotometer (Analytik Jena, Germany). Bromelain concentration was measured using □^1%^_280nm_ = 2.01 **[14].** This preparation is a mixture of bromelain isoforms and minor amount of other cysteine proteases of similar molecular weight **[15].**

### Peptides derived from fruit bromelain under conditions of human digestive system

The fruit bromelain protein pool (10 mg) was dissolved in 2 ml of 5% HCOOH to obtain an acidic solution (pH 2.0) followed by addition of Pepsin (2 mg, 1:50 wt/wt) and incubation at 37°C for 3 – 4 hr. Thereafter, NH_4_HCO_3_ was added to increase pH of the solution to 7.5. Trypsin and chymotrypsin (2 mg each, 1:50 wt/wt) were sequentially added along with trace amount of elastase and carboxypeptidase for further digestion at 37°C for 7 – 8 hr. Incubation conditions were maintained mimicking human gastrointestinal tract digestion.

### Separation of bromelain peptides by Sep-Pak C18 Cartridges

Fruit Bromelain derived peptide pool was separated from undigested protein part by using Waters C18 Sep-Pak Cartridges. Prior to use, cartridges were washed with 10 ml acetonitrile and equilibrated with water containing 0.1% TFA. After sample loading, unabsorbed proteins and large peptides were eluted and cartridge washed with water containing 1% TFA. Peptide pool was eluted with 2 ml 50% acetonitrile containing 0.1% TFA. MS analysis revealed presence of peptides ranging from 200 Da to >1 kDa. For further fractionation according to size, the pool (2 ml) was applied to Sephadex G-10 column (85 ml, fractionation range <700 Da) fitted with Biologic duo flow instrument (Bio-Rad) at a flow rate of 6 ml/hr. Fractions eluted were continuously monitored at 220/280 nm in a UV visible spectrophotometer, collected according to peak positions and further characterized by MS analysis.

### Fluorometric quantification of amyloid aggregates

Th-T assay was employed to follow aggregation kinetics of Aβ. Th-T, a fluorescent dye interacts with β-sheet rich fibrils leading to characteristic increase in fluorescence intensity in the vicinity of 480 nm, relative to unbound Th-T (ex: 450 nm; em: 460-600 nm). λ_max_ emission intensity varies from 480 to 487 nm **[16]**. A Hitachi F4500 fluorescence spectrometer attached to a circulating water bath at 25□ was used (ex/em slit widths 5/5 nm). A stock solution of Th-T (250 μM in water) was made using □_412 nm_ = 35,000 M^−1^cm^−1^ and aliquots of 10 μl were transferred to the reaction mixture before fluorescence measurements were recorded. The reaction mixture consisted of 10 mM Na phosphate, pH 7.5 and 1-10 mM peptide solution in a final volume of 1 ml. Blank correction was done for all (n = 5, with replicates of 5 in each set).

### Determination of Protein and peptide concentration

While bromelain peptide concentration was determined optically using molar extinction coefficient of peptide bond at □_214 nm_ = 923 M^−1^cm^−1^ in presence of acetonitrile and formic acid **[17]**, concentration of Aβ was evaluated using □_276 nm_ = 1450 M^−1^cm^−1^ **[18]**.

### Mass spectrometry analysis

Molecular mass of peptide pool was determined using a Q-Tof Mass Spectrometer (Waters Corporation, USA). The sample was desalted using Zip-Tip μ-C_18_ (Millipore, Billerica, MA, US). Matrix bound samples were eluted in 50% acetonitrile in water containing 0.01% formic acid. Samples were further analyzed under positive mode of ESI at desolvation temperatures of 100-125°C. Argon as a collision gas at 2 kg/cm^2^ having collision energy of 10 eV was applied. Micro channel plate detectors were used.

### Dynamic Light Scattering (DLS)

DLS monitors change in particle size and distribution, and calculates average hydrodynamic radius during protein/peptide aggregation. DLS measurements of different Aβ species in presence and absence of bromelain derived peptides were taken using a back-scatter apparatus (Malvern Nano ZS, Malvern) having a constant scattering angle of 90°, at 25 ± 1°C **[19]**. Samples were diluted to a final concentration of 0.1 mg/ml in a total volume of 1 ml (n = 5, with replicates of 5 in each set).

### Transmission Electron Microscope (TEM)

Samples were placed on a 300-mesh copper grid covered by carbon-coated film and incubated for 10 min at 25°C. The excess fluid was removed and grids were negatively stained for 30 sec with 10 μL of 1% uranyl acetate solution. Excess stain was removed by repeated washing with Milli-Q water and samples were visualized in a TECNAI G2 TEM (Thermo Fischer Scientific, USA) operating at 120 kV accelerating voltage and 1,15,000x. To estimate widths of individual fibres, digital electron micrographs were analyzed by MCID Elite (Micro Computing Imaging Device 7.0, revision 1.0, Imaging Research, Inc.).

### Atomic Force Microscopy (AFM) Imaging

Protein samples (10 μl, 200 ng/ml) were deposited onto freshly cleaved mica sheets and air-dried for 20 min. Sometimes the sample was gently washed with 0.5 ml Milli-Q water to remove molecules that were not firmly attached to the mica and then air-dried as above. Acoustic AC (AAC) mode AFM was performed using a Pico plus 5500 AFM (Agilent Technologies, USA) with a piezoscanner maximum range of 9 μm. The cantilever resonance frequency was 150 – 300 kHz.

Cells were seeded on glass cover slips maintaining a density of 10^6^cells/well in a 6-well plate and treated with media containing 5 μM of Aβ40/42 at 0 hr with or without 5 μM of synthetic peptides, after preincubation for 24 hr. The same media volume was added to control cultures. Cells were incubated for an additional period of 48 hr at 37°C and allowed to reach 70–80% confluency. Cover slips with adherent cells were then washed with 1X PBS to completely remove media, followed by fixation with 1% PFA for 1 hr at 4°C. Prior to imaging PFA was rinsed well with 1X PBS and finally with double-distilled water to prevent deposition of any excess buffer molecule. Imaging was done in dry mode using 100 micron scanner. Cantilevers of 450 μm length with a nominal spring force constant of 0.2 N/m were used. Resonance frequency was set at 13 kHz. Individual plots shown for surface topography of various samples are representative views of morphologies observed for multiple areas of the samples.

In both sets, images (512 by 512 pixels) were processed using Pico scan software (Molecular Imaging Corporation, San Diego, CA).

### Disaggregation of amyloidal protein by bromelain derived peptides

The following experiments were designed to evaluate anti amyloidogenic property of protease digested small peptide pool of fruit bromelain using three-phase study protocol **[20]**: (i) freshly prepared Aβ40 was co-incubated with digested peptide (7 μM) for 96 hr. Aliquots were withdrawn at 0, 6, 20, 72 and 96 hr; (ii) Oligomers of Aβ40 were formed from freshly prepared monomeric peptide upon incubation for 20 hr. Thereafter, aggregation was followed in presence and absence of bromelain derived peptides up to 96 hr. Aliquots were taken at 20, 36, 48, 72 and 96 hr post initiation of oligomerization and, (iii) Matured Aβ40/42 fibrils were formed after incubation of monomeric peptide for 96 hr. Thereafter, bromelain derived peptides were added and disaggregation of fibrils was monitored for 48 hr. In all sets, samples were analyzed by Th T assay and TEM or AFM images. Peptide solutions without aggregates served as control for fluorescence measurements (n = 5, with replicates of 5 in each set).

### Conformational studies

#### Interactions with ANS

ANS, an extrinsic fluorescent probe interacts nonspecifically with surface hydrophobic patches of proteins resulting in significant increase of quantum yield **[21]**. Emission intensity of Aβ (100 μM) was followed in presence of 0 – 500 mM of ANS in 10 mM Na-phosphate, pH 7.5 (ex: 380 nm; em: 400–550 nm; em_max_: 470 nm). Spectral corrections with ANS as control were done for all sets.

#### Fourier-Transform Infrared (FT-IR) Spectroscopy

FT-IR spectra were recorded in a Tensor 27 FT spectrometer equipped with a liquid N_2_-cooled mercury cadmium telluride detector. For each spectrum, water vapor used as blank was subtracted for baseline correction. Spectra were obtained using Bruker, Opus Software and scans were taken between 1,590 and 1,710 cm^−1^ and normalized to unity.

#### Circular Dichroism (CD) spectroscopy

CD spectra of Aβ (30 ng/ml) were analysed using a Jasco J-815 spectropolarimeter (JASCO International, Japan). A cell having 0.1 mm optical path length and a bandwidth of 1 nm was used. Solvent spectra were subtracted from the measured spectra in each experimental set. All spectra were recorded in the far UV range of 195 – 250 nm at 25°C considering an average of ten scans. Spectral analysis was done with Origin Lab 8.0 software.

For all sets n = 5, with replicates of 5 in each set were maintained.

### *Ex-vivo* studies

#### Cell culture

PC 12 cells were purchased from American Tissue Type Collection (ATCC), Virginia, USA. Cells were cultured in DMEM containing 10% FBS and 1X penicillin and streptomycin mixture at 37°C in a humidified incubator having 5% CO_2_ environment. Cells were seeded and allowed to reach 80 – 85% confluency before performing experiments. Five experimental sets were maintained: (1) untreated cells; (2) cells incubated with Aβ peptide (10 μM final concentration) for 24 hr; (3) cells treated with bromelain peptide (25 μM final concentration) for 48 hr; (4) cells treated with Aβ for 24 hr followed by bromelain for 48 hr and (5) cells co-incubated with Aβ and bromelain for 48 hr. Around 10^5^ cells/100 μl of medium/well were maintained in a 96 well polystyrene plate. Cell viability was quantified for all sets using 3-(4,5-dimethylthiazol-2-yl)-2,5-diphenyltetrazolium bromide (MTT) assay after the incubation period.

#### Cell viability test (MTT assay)

Mitochondrial respiration, an indicator of cell viability, was assessed in experimental sets (n = 4), using the mitochondrial dependent reduction of MTT to formazan **[22].** MTT (10 μl from a stock of 5 mg/ml in PBS was added to each well and incubated for 4 hr at 37°C. The medium was then aspirated and replaced by 100 μl of DMSO, following which plates were agitated at 25°C for 10 min and absorbance recorded at 590 nm by a multi well plate reader (Biotek–Epoch; Biotech Instruments, Winooski, VT, USA). The average absorbance value of replicate wells was considered for each set. In these experiments, cells without test samples but MTT served as positive control (blank) while cells treated with hydrogen peroxide (10 μl) served as negative control.

### In *vivo* studies

#### Experimental Animals

Adult male Sprague-Dawley rats weighing 280 – 330 g were procured from the random bred colony of the animal care facility of the institute (IICB) and were maintained following good husbandry conditions at standard temperature (24 ± 4°C), humidity (60 ± 5%) and 12 hr light-dark diurnal cycles. They were provided with food and water *ad libitum*. All animal experiments were carried out as per guidelines of Institutional Animal Ethics Committee for the Purpose of Control and Supervision of Experimentation on Animals (CPCSEA), under the Division of Animal Welfare of the Ministry of Environments, Forests & Climate Change, Government of India.

#### Brain Stereotaxic Surgery

Rats were anesthetized intraperitoneally with pentobarbital (50 mg/kg i.p.; Thiosol, Neon Laboratories Limited, Mumbai, India). Animals were fixed in a stereotaxic apparatus (Stoelting, MO, USA) according to Paxinos and Watson (1998) with the incision bar kept 3.5 mm below the interaural line and their body temperature maintained at 37°C **[23]**. We initially compared various doses of both Aβ and bromelain to analyse the dose dependency of bilateral intracerebroventricular (ICV) infusion of both in our hands, and thereby optimize the dose (Aβ, 6 μM/ rat and bromelain, 20 μM/ rat) for developing animal model in our laboratory. The stereotaxic coordinates used were: Lateral = 0.12 cm, Anterio-posterior = −0.90 cm and Dorsoventral = 0.34 cm, with reference to Bregma point following the Rat Brain Atlas of Paxinos and Watson **[24]**. In the control group, artificial Cerebrospinal fluid (aCSF) (147 mM NaCl; 2.9 mM KCl; 1.6 mM MgCl_2_; 1.7 mM CaCl_2_ and 2.2 mM dextrose) was infused ICV (3 μL into each side). All infusions were done bilaterally with each hemisphere at a time in a total volume of 12 μL (6 μL/side) at a flow rate of 0.5 μL/min. The infusion probe was left in position for an additional five minutes after each drug delivery for proper diffusion of drug into the ventricles. Proper postoperative care, including hand-held feeding was provided until animals recovered completely.

Experimental sets comprised of animals treated with: (1) Aβ, (2) bromelain, (3) Aβ followed by bromelain after 21 days and (4) Aβ+-bromelain simultaneously. On the respective day of sacrifice (21 days post infusion of Aβ and 14 days post infusion of bromelain), the rats were anaesthetized with urethane and transcardially perfused with 1X PBS (pH 7.2) followed by 4% paraformaldehyde in PBS for 5 min each,. The brains were removed under perfusion conditions and fixed overnight in PBS containing 4% paraformaldehyde and subsequently transferred into 30% sucrose in PBS after 48 hr to allow cryoprotection.

### Histology

Brain samples stored in PBS–30% sucrose solution was investigated later by tissue histopathology using hematoxylin-eosin staining. Slides were prepared following usual H&E staining post parafilm block preparation and tissue sectioning using manual microtome machine (Leica RM2235 Manual Rotary Microtome for Routine Sectioning). Histological analysis of wound tissue samples was carried out with an Olympus SZX 10.

### Statistical analyses

All experimental results have been reported as mean ± SEM. Student’s t-test was performed in each case to evaluate significant difference between means and has been represented as p values. Number of replicates performed for each experiment has been mentioned in the respective sections.

## RESULTS

### Destabilization of Aβ amyloid by bromelain derived peptides

Preliminary experiments with proteolytically active ‘fruit bromelain’ indicated that the enzyme was capable of destabilizing preformed Aβ40/42 fibrillar structures to small oligomers. This was also observed using inactive or proteolytically degraded ‘fruit bromelain’ or even synthetic peptides using specific template of ‘fruit bromelain’ sequence. Therefore the underlying mechanisms might be proteolysis or interaction between amyloid structures and specific amino acid stretches of fruit bromelain or a combination thereof. Mass spectrometric analysis of dissociated products of Aβ amyloid never revealed any fragment smaller than the monomeric Aβ peptide.

The pool of eluted peptides were capable of disaggregating preformed Aβ40 (10 μM) in a concentration dependent manner (0 – 40 μM) as suggested from ThT assay. Under experimental conditions, approximately 70% of disaggregation was achieved after 24 hr of incubation (Fig. 1A). AFM images demonstrated dense fibrillar structure of pretreated Aβ amyloid aggregates (Fig. 1B, upper panel, left) while they followed course of degradation in presence of bromelain peptides (Fig. 1B, lower panel, left). During imaging, the skeleton of fibril though visible, at places the connections were loose. Fibril structure of β-amyloid aggregate was degraded by peptide pool, compared to matured fibrils of control sample. Corresponding TEM micrographs showed that fibrillar network was composed of rod-like structures of variable length and diameter that were fragmented to small spherical oligomers to monomer like molecules (Fig. 1C, upper and lower panels). As a negative control, it was ensured from both AFM and TEM that the fibrillar structure was stable under conditions of incubation with peptides for 24 hr. Encouraged by this observation, peptides of Mw <500 Da were isolated from bromelain digest as they may enter into the blood stream, Mw being one of the precondition to cross BBB.

**Fig. 1:**
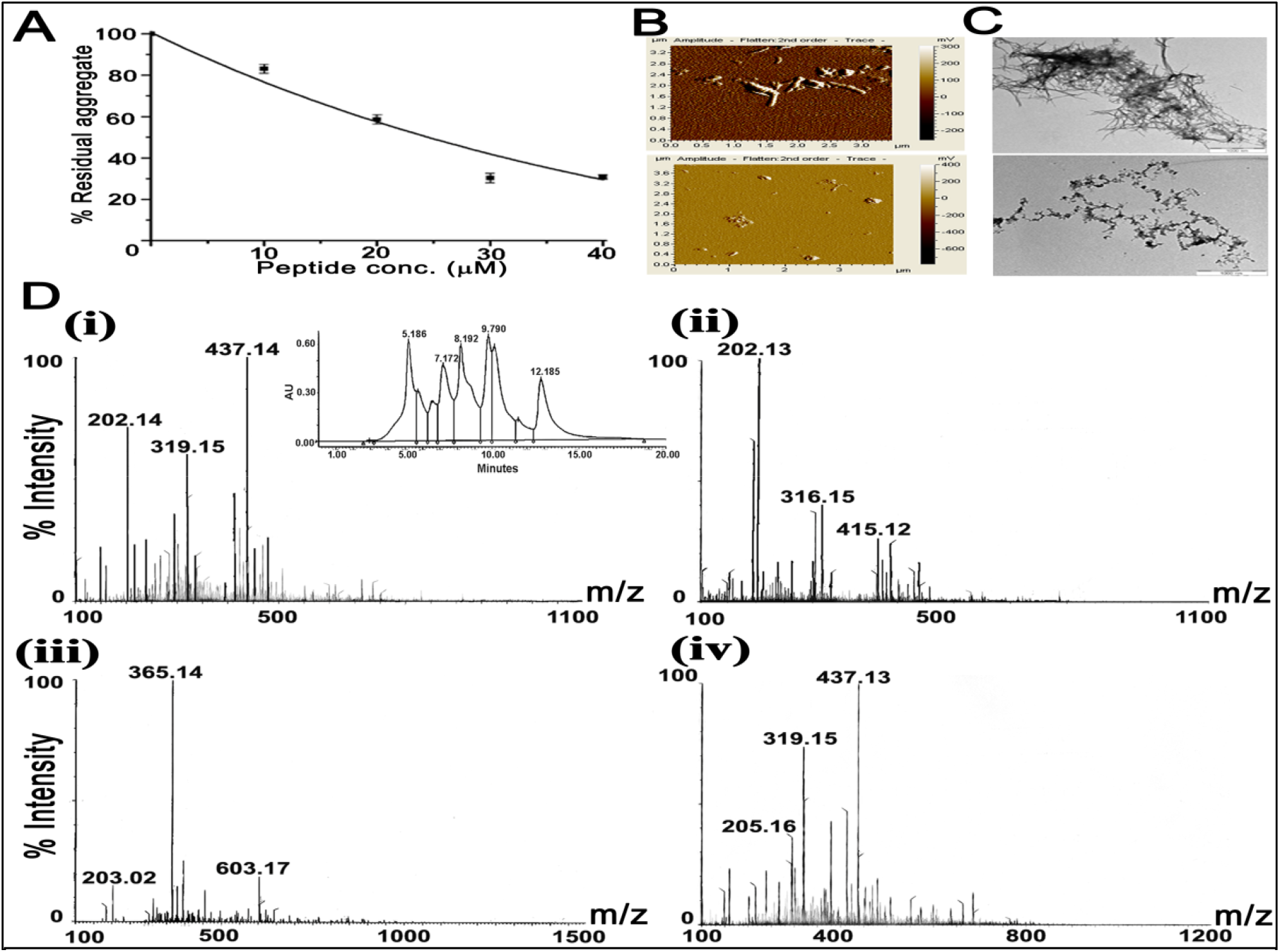
Destabilization of preformed Aβ40 aggregates and ESI mass analysis of different fractions of peptides. (A) Dependency of destabilization of Aβ 40 aggregates on concentration of peptides. Residual structure was measured by Thioflavin-T assay. (B) AFM image of the preformed Aβ40 aggregate of 72 hr (upper panel) and the disaggregated state (lower panel). (C) Corresponding TEM images have been shown in (upper panel) and (lower panel). Experimental conditions have been described in the text. (D) Fruit bromelain was treated with digestive enzymes and the peptide pool was separated using Sephadex G-10 size exclusion column. The peak fractions corresponding to retention times (R_t_) 5.186, 7.172, 8.192 and 9.790 min were analyzed by ESI-MS (i-iv). Being in the desalting zone, the last chromatographic fraction of R_t_ = 12.185 min was not analyzed. The HPLC profile has been shown in Inset (D(i)). See also Fig. S1.

### Isolation of bromelain derived peptides of Mw <500 Da

SE-HPLC profile of peptide pool generated from freshly prepared pineapple after human protease digestion (Fig. S-1A) revealed unresolved overlapping broad peak. 15% SDS-PAGE indicated diffused band around 10 kDa (Fig. S-1B). Corresponding MALDI-MS analysis also deciphered mixture of undigested protein part and digested polypeptides ranging from 1000–5000 Da (Fig. S-1C). Most of these polypeptides were >700 Da, the molecular mass limit for a substance to cross BBB **[25]**. The viability of peptide-based aggregation mediator is determined by its ability to cross BBB and withstand *in vivo* conditions leading to mediator degradation. Low molecular weights are needed for crossing BBB and the transport must also be done without any degradation of aggregation mediator **[26]**.

The chromatogram from Sephadex G -10 indicated presence of more than 10 components (Fig 1D). The Mw of the 4 trailing fractions were verified by MS analysis (Fig. 1D, marked as (i) – (iv) Inset). Except one peptide of Mw 603.17, Mw of all major peptides of these 4 fractions were <500 Da (200-437 Da) (Fig. 1D). These fractions were pooled as stock solution for subsequent experiments. Fractions eluting ahead of Fraction I with Mw >500 Da were excluded.

### Destabilization of Aβ 40 aggregate by digested bromelain derived small peptides of Mw <500 Da

#### Th T assay

Disassembly of preformed Aβ40 fibrils by peptide pool was both concentration and time dependent. With increasing peptide concentration from 0 – 40 μM keeping duration of incubation and temperature at 24 hr and 37°C respectively, aggregates exponentially destabilized having a residual structure of around 20% (Fig. 2A). Similarly, increasing incubation time from 0–12 days keeping peptide concentration and temperature at 7 μM and 37°C respectively, an exponential time course having 85% disaggregation was observed (Fig. 2B).

**Fig 2:**
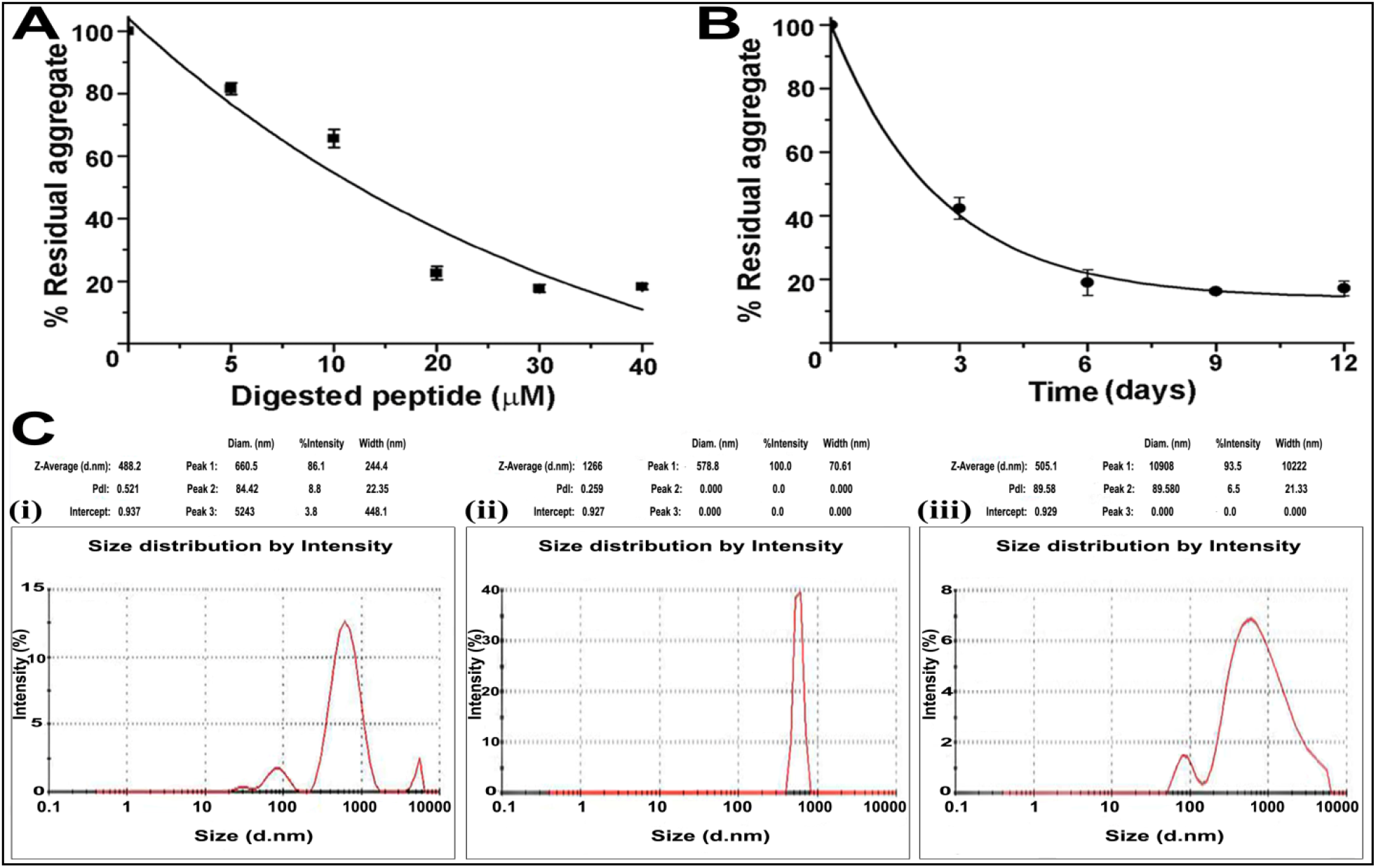
Destabilization of Aβ40 aggregate and analysis of hydrodynamic radius of different forms of Aβ40. Dependency of disaggregation on (A) the concentration of pool of digested peptides (1-40 μM) in 48 hr and (B) time using 7 μM of the peptide pool as measured by Th-T fluoremetric analysis. (C) DLS measurements of (i) Monomeric Aβ40, (ii) Aβ40 fibrils and (iii) preformed Aβ40 aggregates after incubation in the presence of fruit bromelain peptides after 48 hr of incubation at 37°C.

#### DLS

DLS profiles of monomeric, aggregated and disaggregated states of Aβ showed actual particle distribution according to size in diameter avoiding stability factors against shearing forces (Fig. 2C). Average diameter of particles present in these sets were 488.2 nm, 1266 nm, 509.1 nm for monomer, aggregate and disaggregate respectively as observed within scale of detection. In case of aggregated state, majority of the particles were out of scale and could not be detected. The correlogram coefficient data of DLS shows the time at which the correlation starts to significantly decay as an indication of mean size of the sample. Further, high monodispersity of a sample is a measure of how steep the line is and conversely, polydispersity of the sample is indicated by how extended the decay takes (figure not shown). These features were in good agreement with DLS profiles (Fig. 2C). Th-T and DLS analysis preliminary indicate Aβ disaggregation potency of peptide pool.

### Inhibition of Aβ 40 aggregate formation by digested peptide pool

#### Digested peptide pool inhibits Aβ40 aggregate formation from monomers

Time course of Aβ40 aggregate formation in presence and absence of bromelain peptides was followed for 96 hr. TEM images revealed that even at 0 hr, monomeric Aβ 40 was contaminated with trace amounts of very small multimeric components. Samples represented in Fig. 3A are electron micrographs reveal clear time dependent aggregation kinetics at 0, 6, 20, 72 and 96 hr time intervals. While oligomerisation was noticed after first 20 hr, aggregates were visible roughly around 48 hr onwards. But in the presence of digested peptide, there were no aggregates even up to 96 hr indicating total abolition of aggregation formation. All three phases of Aβ40 aggregation kinetics was monitored through Th-T fluorescence at same time intervals (0 – 96 hr) (Fig. 3C) exhibiting a sigmoidal curve showing static fluorescence intensity up to 20 hr, indicating lag period represented by presence of monomers and small oligomers. After which, there was an intermediate phase from where larger oligomers and other intermediate forms might have formed till 48 hr followed by the last saturation phase where, fully matured fibrils are formed thereafter, where Th-T fluorescence became static. In presence of digested peptide, this sigmoidal pattern of Aβ40 aggregation kinetics was disturbed, and showed only a lag phase with monomers or small oligomers only. This has been supported by AFM images (Fig. S-2A).

**Fig 3:**
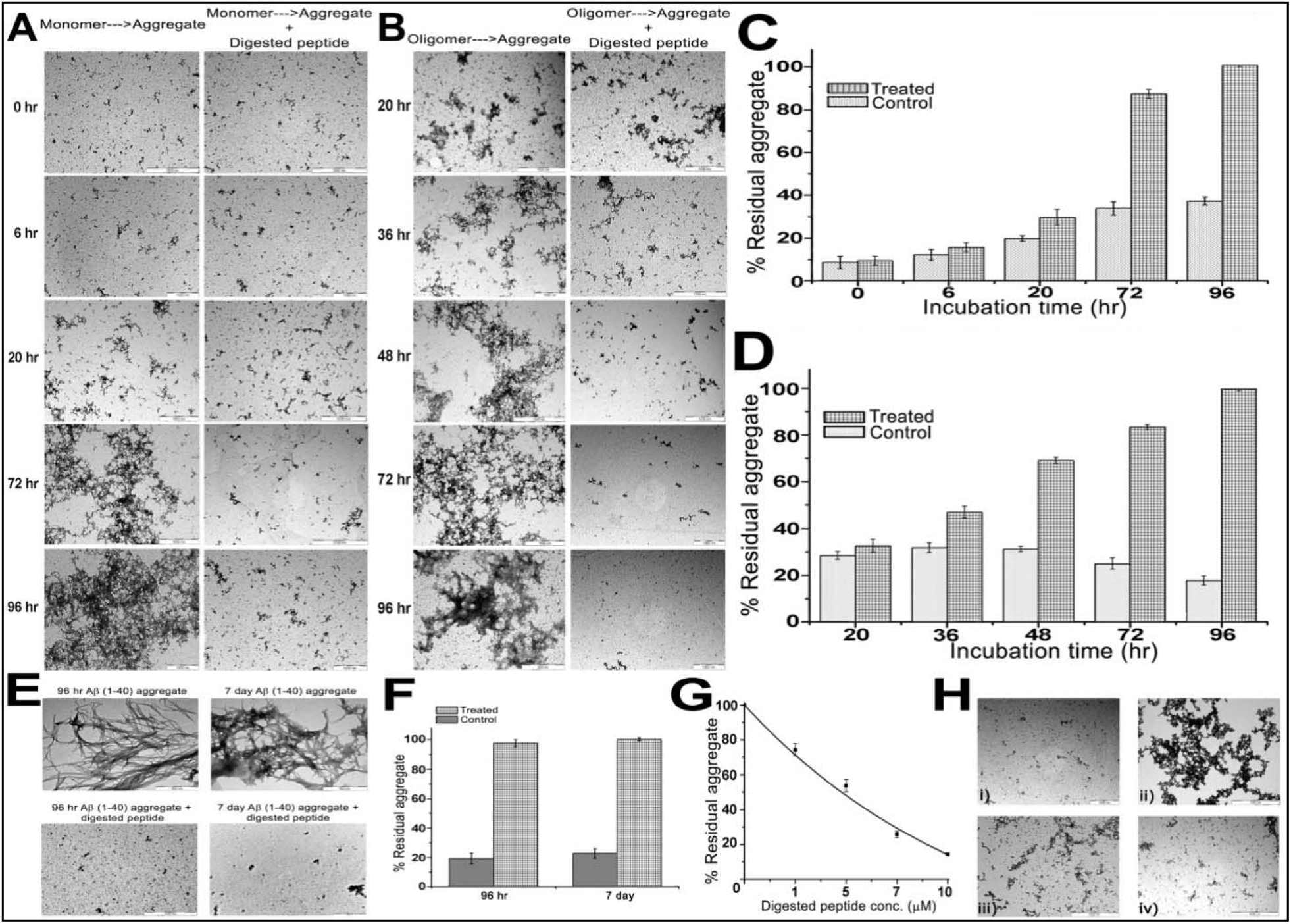
Aggregation kinetics of Aβ40/42 fibrils. (A) Monomer to aggregate: Aβ40 was incubated in absence (left panel) and presence (right panel) of 7 μM digested peptide at 37°C. (B) Oligomer to aggregate: Oligomers were generated after incubating Aβ40 for 20 hr under fibrillating conditions were incubated under similar conditions as in A. Aliquots were withdrawn at time intervals as indicated vertically. Corresponding Th graphs of A and B have been depicted in C and D respectively. In both panels, emission intensity of the sets at 96 hr was considered as 100% (See also Fig. S2). (E) TEM images: (Upper panel) Aβ40 incubated under defined conditions of fibrillation for 96 hr and 7 days. (Lower panel) Corresponding sets after incubation with peptide pool for 7 days. (F) Th T assay of the four samples presented in E where absence and presence of the peptides have been denoted as control and treated. Intensity of the control sample after 7 days has been considered as 100%. (G) Estimation of concentration of peptide pool as measured from Th T assay during disaggregation. Fluorescence from spontaneously formed aggregates after 7 day was considered as 100%. (H) TEM images of (i) monomeric Aβ42, (ii) monomeric Aβ42 fibrillation for 96 hr at 37°C; (iii) fibrillation as seen in presence of peptide pool for 48 hr and (iv) protease digested peptide pool, inhibiting Aβ42 aggregation.

#### Digested peptide pool inhibits Aβ¦40 aggregate formation from oligomers

Fig. 3D shows Th-T fluorescence data of Aβ40 in the presence and absence of digested peptide pool from the oligomeric stage onwards at 20, 36, 48, 72 and 96 hr time intervals. Th-T fluorescence increased from 20 to 96 hr indicating formation of fibrils from oligomers stage on contrast to a significant decrease in the intensity profile in presence of peptides, indicating inhibitory effect on oligomers even after aggregation continued up to 20 hr. These results were further confirmed by TEM (Fig. 3B) and AFM images (Fig. S-2B).

#### Digested peptide pool disaggregated preformed Aβ40 fibrils

Under defined conditions of self-aggregation, Aβ40 was incubated and aliquots withdrawn for TEM imaging. The sample after 96 hr of incubation showed an extensive network of branched fibrillar structures having high degree of cross-links. With increase of incubation period up to 7 days, overlapping fibrillar structures became denser and more matured with heavy branching. In either set, no trace of oligomeric components could be detected. (Fig. 3E upper panel). When these samples were separately treated with bromelain peptides for 7 days at 37°C, dense fibrillar structures completely dissociated into small oligomers and further down to monomer or dimer that remained undetected by TEM. Very small pieces of broken fibrillar structures were visible in the 7 days sample as compared to 96 hr (Fig. 3E lower panel). Samples were also treated with Th-T to estimate presence of aggregated structures. Assuming that the emission intensity of the self-aggregated sample after 7 days was 100%, the 96 hr incubation sample offered nearly 95% emission suggesting complete aggregation. In either set, disaggregated fibrils offered nearly 20% of emission intensity. Considering that Aβ40 in its monomeric condition at 0 hr shows nearly 10% emission, Th-T assay indicates that disaggregation was nearly complete in each set (Fig. 3F).

### Disassembly of pre-formed Aβ42 fibrils by digested bromelain peptides

The potency of Aβ42 to form amyloid aggregate is higher than Aβ40 and therefore, is clinically more important. Though the ‘sticky’ hydrophobic regions of the two peptides (15 – 21) are identical, presence of two additional residues at C-terminal end of Aβ42 renders it more toxic **[27]**. TEM image of Aβ42 monomers showed presence of trace amount of small oligomers as physical impurity (Fig. 3H(i)). The monomer on self-aggregation for 7 days under defined conditions exhibited dense overlapping fibrillar network with extensive branching where oligomers remained undetected (Fig. 3H(ii)). Preformed Aβ42 fibrils when treated with bromelain peptides, defibrillated completely within 7 days. At an intermediate time course of 4 days, dissociation of network structure was observed with generation of varied oligomers where links between large oligomers were either nonexistent or very feeble (Fig. 3H(iii)). With passage of time, moderately large oligomers reduced in size and links between them were abolished (Fig. 3H (iv)). Defibrillization was dependent on peptide concentration (0 – 10 μM) holding all other experimental conditions constant as observed from Th-T. Acknowledging limitations of Th-T and interference from monomeric Aβ42 peptides, it may be stated that bromelain peptides (10 μM) can efficiently dissociate preformed Aβ42 fibrils into small oligomers (Fig. 3G).

### Change in secondary and tertiary structures

To have an insight of the molecular structure of Aβ40 peptides during the course of aggregation and disaggregation, interaction with the fluorophore 8-ANS sensing hydrophobic patches of anchoring proteins, FT-IR and CD spectroscopy were applied. Four states of Aβ40 were characterized; the monomeric state, self-aggregated state where fibrillar structure was formed by 7 days, the same incubate in presence of bromelain peptides where aggregation was inhibited and disaggregated state from preformed fibrils after interaction with bromelain peptides.

#### Interaction with ANS

Interaction of aforementioned states of Aβ40 with 0 – 300 μM ANS was followed. It showed significant and comparable interactions with monomeric, aggregation inhibited and disaggregated states where the interaction with aggregated state was greatly reduced. In all sets, emission intensity reached plateau level between 450 – 500 μM of ANS indicating saturation of ligand binding. Considering emission intensity of disaggregated state as 100%, relative emissions from monomeric, aggregation inhibited and aggregated states were 95.54, 90.27 and 45.99 % respectively (Fig. 4A). This profile also shows that structures of the three sample sets constituting monomer and small oligomers were similar but not identical so far their interactions with ANS was concerned. A simplified explanation for low ANS binding with aggregated state is that hydrophobic stretches of constituent molecules were no more available. To ensure that low affinity of ANS with aggregated state did not arise from an artifact, time course of aggregate formation was followed in presence of 500 μM of ANS up to 96 hr. Dduring this period, reduction of emission intensity followed an exponential pattern leading to 64% while considering the emission at 0 hr as 100%. (Fig. 4B). The dissociation constant (K_d_) of the conjugate has been calculated to be 11.171 μM.

**Fig 4:**
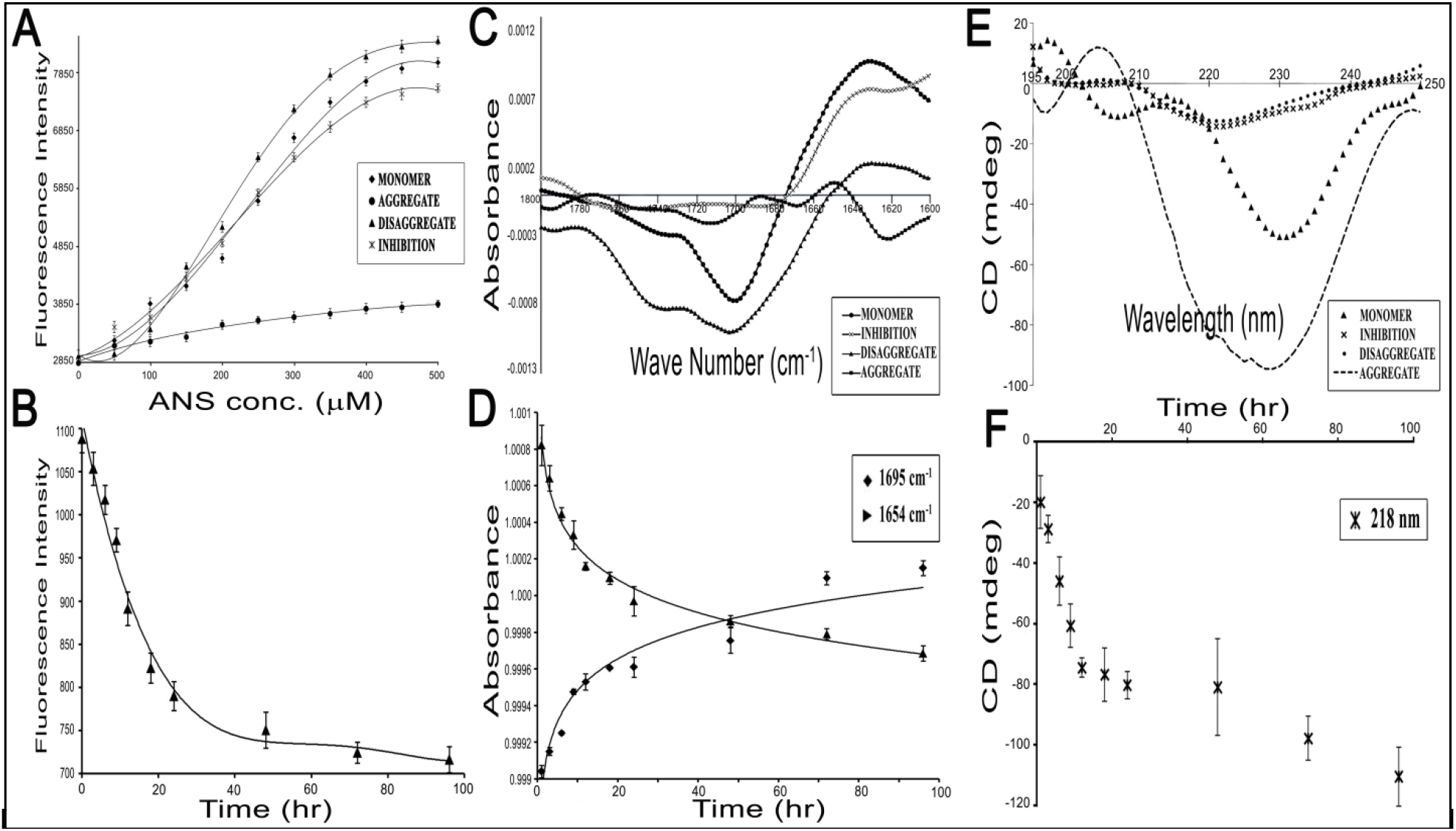
Spectroscopic analysis of different forms of Aβ40. (A) Concentration dependency of ANS interaction as observed from fluorescence emission intensities. (B) Interaction of ANS (500 μM) as followed with different species of Aβ during the time course of aggregation. (C) FT-IR spectra ranging from 1800-1600 cm^−1^ for different species of Aβ. (D) Change of secondary structure of Aβ40 during fibrilization was followed at 1695 cm ^−1^ and 1654 cm^−1^. (E) Far UV CD spectra (195–250 nm) of these states and (F) Time course of aggregate formation from monomeric state as observed from ellipticity values at 218 nm. Each spectral value is the average of 5 runs and in all sets, buffer corrections have been done. The descriptions of the notations have been provided in respective insets. All experimental conditions have been mentioned in details in the text.

#### FT-IR

Change in protein secondary structure could be detected from FT-IR spectra within 1800 – 1600 cm^−1^. While bands in the range of 1648 – 1657 cm^−1^ are assigned for α-helix, those between 1623 – 1641 cm^−1^ and 1674 – 1695 cm^−1^ are assigned for high frequency β-sheet components. Since rotational-vibrational spectra of proteins are very sensitive to their electronic environment, no single wave number could be assigned to these protein structures **[28].** FT-IR spectra of monomeric, aggregated and disaggregated Aβ40 peptide and the inhibited form were scanned in along 1590 – 1710 cm^−1^. Significant differences of spectra and therefore structures were predicted (Fig. 4C). While correlating these results with structure, significant loss of residual α-helices and gain of β-sheet structure of monomeric Aβ40 during aggregate formation was observed between 0 – 96 hr at 1654 cm^−1^ and 1695 cm^−1^ respectively **[29]** (Fig. 4D). FT-IR results though semi-quantitative, characteristic features of protein aggregate formation were evident.

#### CD

Far-UV CD spectra of aforementioned states between 195 – 250 nm showed distinct change of secondary structures (Fig. 4E). β-sheet content of monomeric peptide was calculated to be 24% that was comparable to 27% as reported **[30]**. This minor difference is expected to arise from solvent composition initially used to solubilize the peptide. An increase of negative ellipticity of aggregated state was evident indicating rigidity of structure. The disaggregated state and the peptide resisting aggregation showed similar low ellipticities in the whole spectral range indicating generation of distinctly different conformers and prevalence of random coil structures. To follow formation of aggregated state from monomers through oligomeric states, change of ellipticity was followed for 96 hr at 218 nm. A continuous decrease of negative ellipticity was indicative of β-sheet formation, a characteristic feature of amyloid-like structures. The decrease appeared to be continuous as fibrillar structures require longer time for maturation (Fig. 4F). It is noteworthy that the disaggregated state and monomer that inhibits aggregate formation had very similar random coil rich structures.

#### Sequence alignments of fruit bromelain and Aβ peptide using ClustalW2

Amino acid sequence of fruit bromelain constituting 351 amino acids (UniProtKB-O23791) and Aβ40/42 were aligned using ClustalW2 (multiple sequence alignment) software (Fig. 5A). Previous studies reported ^16^KLVFFAE^22^ of Aβ40/42 as the most aggregate prone zone of the peptide that forms the core of aggregate from which fibrils propagate **[31]**. Reports assign tryptophan as a potent residue for Aβ fibrillation and plaque formation **[32]**. Alignment of sequences of fruit bromelain and Aβ40/42 peptides showed significant degree of homology around the aggregate prone zone of Aβ peptide. Alignment of fruit bromelain to Phenylalanine rich sequence KLVFFAE of Aβ40 [Clustal W 2.0] and analyses of peptides generated after gastrointestinal digestion deciphered [Expasy Peptide Cutter] probable bromelain peptides as TIIGY and GQD. In vitro experiments conducted with these peptides indicated specificity of small bromelain peptides in disaggregation as visualized from microscopic images (Fig. 5B) and was hence further used for *in-vivo* and *ex-vivo* analyses.

**Fig 5:**
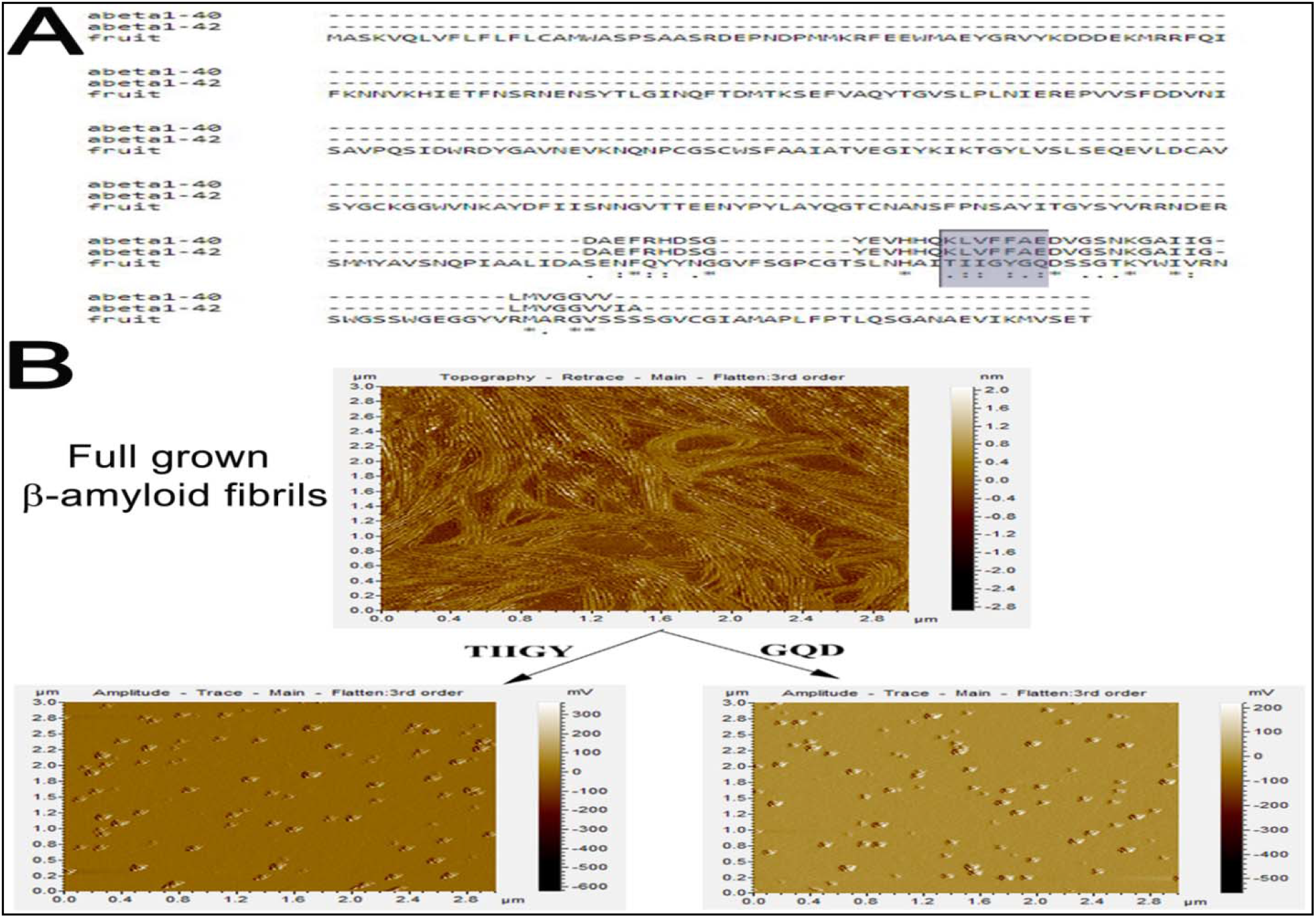
Determination of small peptides corresponding to Aβ. (A) Sequence alignment of fruit bromelain with Aβ40/42 using ClustalW2 software. The aggregation prone region of Aβ40/42 (KLVFFAE, residues 16-22) has been highlighted to denote the corresponding region in fruit bromelain. Two peptides (TIIGY and GQD) corresponding to the highlighted region of fruit bromelain were considered for further studies. The symbols ‘*’, ‘:’ and ‘.’ indicate identical, highly similar and similar residues respectively. (B) AFM images of preformed Aβ40 fibrillar structure (upper panel) and as obtained after incubation for 96 hr with the synthetic peptides TIIGY (lower left panel) and GQD (lower right panel).

### Cellular Studies

#### Cell viability assay

Based on positive responses of *in vitro* experiments, viability of PC12 cells was checked based on dose dependency to optimize Aβ and bromelain peptide concentrations suited non-toxic to cells. The measured cytotoxicity increased significantly with increasing concentrations of Aβ40 within the range of 20 μM – 20 mM. An optimum concentration of 7 mM was optimized for further experiments. This concentration though higher than physiological conditions was maintained to enhance rate of reaction within experimental timeframe (Fig. 6).

**Fig 6:**
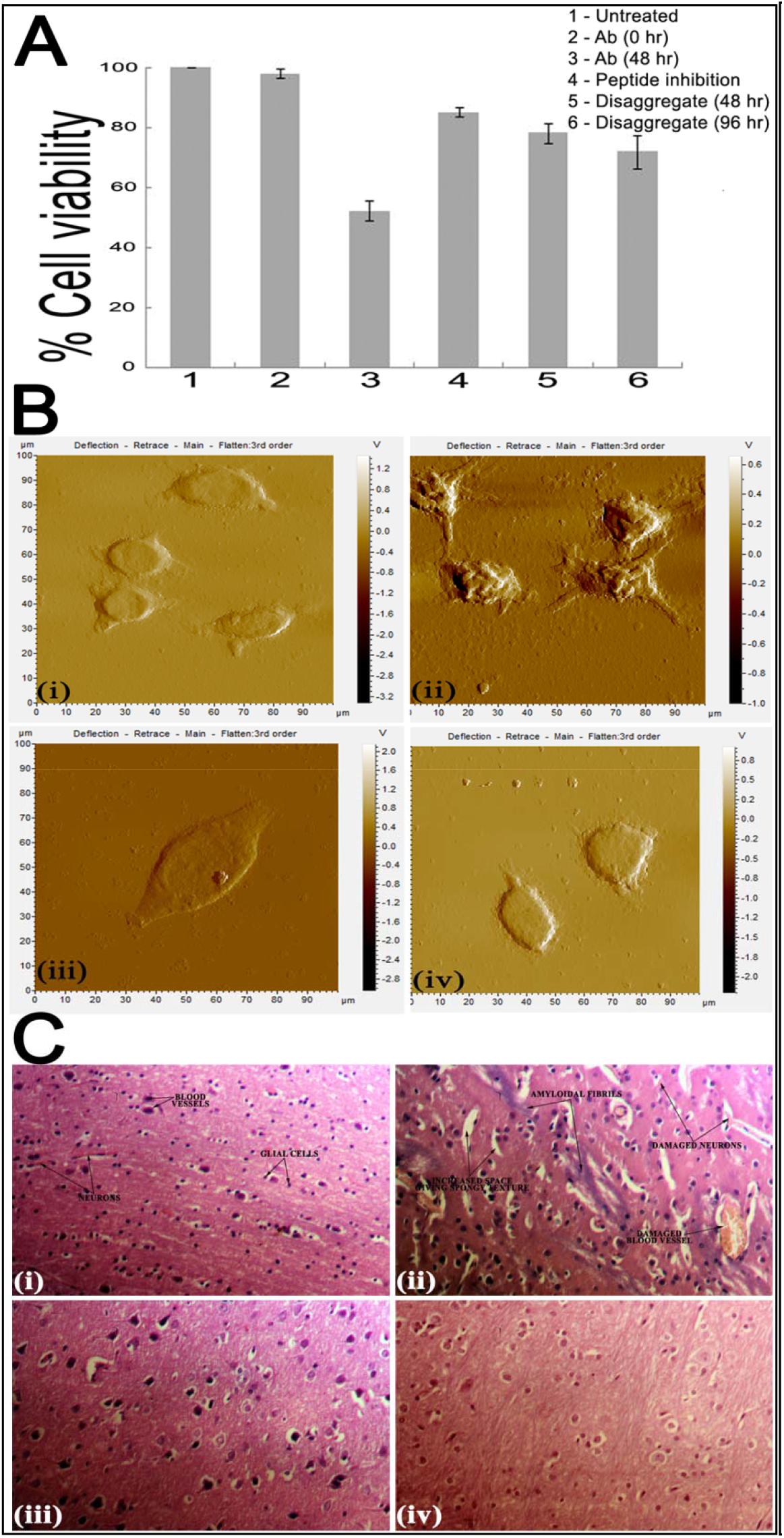
Ex-vivo and in-vivo toxicity studies of Aβ40. (A): Inhibition of Aβ40 induced cytotoxicity by bromelain derived peptides on PC12 cells as observed from MTT assay. Viability of cells has been presented with respect to untreated cells as 100%. All results have been presented after blank corrections where no cell was added. Added reagents have been mentioned in the inset. (B) AFM images of PC12 cells, (i) untreated cells, (ii) cells after 48 hr of Aβ40 treatment. (iii) cells as of (ii) after an additional incubation of 48 hr with bromelain derived peptides and (iv) cells were treated with Aβ40 and peptides. (C) Histological H & E stained section of rat brain tissue obtained from stereotaxic experiments of (i) untreated (control) (ii) Aβ treated; (iii) Aβ treated rats infused with bromelain peptides after aggregation was standardized and (iv) rats treated simultaneously with Aβ and bromelain derived peptides.

Viability of untreated cells in DMEM medium at 0 and 48 hr of incubation was indistinguishable as quantified by MTT assay and was considered as 100%. Viability of cells after treatment with Aβ at 0 and 48 hr was 99.01 ± 1.5% and 52.23 ± 3.4% respectively (Fig. 6A, lanes 1, 2 and 3). Cells when co-incubated with Aβ and bromelain peptides for 48 hr, exhibited viability of 85.04 ± 1.5% [lane 4], when treated with peptides following Aβ deposition exhibited 78.21 ± 3.3% and 71.91 ± 5.6% viability after 48 and 96 hr respectively. On the contrary, cells left for an extended period of 2 – 3 days with Aβ without bromelain peptide treatment gradually lost viability and died. Considering the stress pre-imposed on cells by Aβ deposition, toxicity exerted thereafter by the peptides was insignificant as cells treated as control with only bromelain peptides showed viability of 98.26 ± 3.7%. Under identical conditions, viability of cells treated with hydrogen peroxide was 2.16 ± 1.6%. These results ensured that under conditions of peptide treatment, toxicity was removed and cells propagated (Fig. 6A).

#### Microscopic analysis of aggregation and destabilization on cell surface

PC12 cell topography as visualized by AFM concealed aggregate-like deposits on cell surface upon incubation with preformed Aβ40 monomers (10 μM) 72 hr (Fig. 6B(ii)) on contrast to clear surface of naïve untreated cells (Fig. 6B(i)). The cell morphology also showed a distinct change with irregular shape, membrane distortions and loss of dendritic growth compared to control cells exhibiting typical elongated structure with well-defined dendrites. While cells treated with Aβ and bromelain peptides simultaneously (Fig. 6B(iii)) showed regular shape and dendritic processes, those treated with bromelain peptides after Aβ deposition (Fig. 6B(iv)) was indicative of gradual recovery from stressed conditions illustrating smaller dendritic processes. Therefore, destabilization of preformed fibrils and inhibition of deposition by synthetic peptides was specific.

### Animal Studies

Animals infused with Aβ stereotaxically underwent gradual, significant loss in body weight with time (up to 185 – 220 g), whereas control rats that received only aCSF and those that received an infusion of Aβ along with bromelain together maintained normal gain in body weight (280 – 330 g) during the same time. A similar observation was also noticed for rats that received bromelain peptides after 21 days post Aβ infusion that initially lost weight but with treatment of bromelain peptides over a span of 14 days regained weight to a range of 250 – 300 g, in contrast to those treated with Aβ and bromelain peptides simultaneously that did not show any significant change.

Noticed behavioral changes included decrease in water and food consumption in rats under stress conditions of Aβ treatment only. Behavioral changes showing decreased activity and gradual change of body mass were supported with histological images obtained from H & E staining of brain tissues of corresponding rats. Sham-controlled rats and those that received aCSF infusion showed clear cortical region with homogeneously spaced brain lobes (Fig. 6C(i)), while those infused with Aβ exhibited a spongy appearance, brain necrosis, marked presence of plaque formation in cortical areas of brain and also exhibited cerebral lobes that were much more spaced with increased intraventricular area. Some of the nuclei showed a ring appearance (Fig. 6C(ii)). Brain samples of both, rats infused with Aβ and bromelain peptides simultaneously from day 0 (Fig. 6C(iii)) and AD infused rats treated with bromelain peptides after 21 days (Fig. 6C(iv)) gave images having an improvement in histopathological changes compared to that of AD control but similar to control rat brains (Fig. 6C).

## DISCUSSION

In a series of studies, we demonstrated that amyloid aggregates of Aβ peptide and insulin **[33]** could be destabilized by fruit bromelain peptides. In these sets, synthetic peptides derived using template of proteins was independently capable of destabilizing amyloid aggregates. Specificity of these peptides was evident when similar peptides showed inefficiency in performing the same. A common feature in these combinations is that, presence of intact and functional protein or enzyme was not essential to cause destabilization. Proteolytically degraded enzymes and/or even fragments of intact molecules were also capable of disrupting fibrillar structures. This clearly showed that proteolysis was not involved in the dissociation process. Observations further reinforced that fibrillar structures never produced fragments below the Mw of monomeric Aβ peptide or insulin **[33]**. This was confirmed from mass spectrometric analysis.

Success of these *in vitro* experiments does not qualify these peptides causing defibrilization – no matter whether in a pool or purified or synthetic, to act in brain cortex where Aβ fibrils are formed. Major concern is whether they can cross the BBB **[34]**. Several criteria of the peptides need to be fulfilled to cross BBB viz., hydrophobicity, low molecular weight, high degree of lipid solubility, charge residues of peptides etc **[25]** of which Mw is a fundamental criterion. Under physiological conditions, peptides of Mw more than 400 – 600 Da are generally excluded by BBB **[35]**. Though this stringency is not strictly maintained in case of AD patients where compactness of brain cells is affected allowing relatively bigger molecules to enter, in this study, peptides of Mw <500 Da were separated from undigested and large peptides of fruit bromelain after extensive protease digestion followed by gel filtration and their Mws were verified from MS analysis (Fig. 1D). Based on Clustal W alignment and ExPasy software (Peptide Cutter, using peptides pepsin, trypsin, chymotrypsin, carboxypeptidase and elastase), two peptides TIIGY and GQD were synthesized. Their Mws were 565.67 and 318.29 Da and hydrophobicities were 6.6 and −7.4 respectively **[36].** The hydrophilic peptide was included because of its uncertainty of functioning with respect to BBB under diseased conditions. These peptides not only destabilized Aβ fibrils *in vitro* but also protected PC12 cells against death from assault of Aβ sedimentation (Fig. 6A). These *in vitro* and *ex-vivo* information were essential to initiate animal model experiments (Fig 6B & 6C).

Since pineapple is a widely consumed fruit, efficacy of peptides produced from fruit bromelain after human digestive conditions were verified in rat model. This is strong evidence that if peptides get access to Aβ fibrils in brain, probably they could dissociate the deposits. In this regard, an experiment demonstrating passage of these peptides through artificial brain cell barrier is welcome. Such experiments are performed with synthetic peptides tagged with a small amount of a positron-emitting radioactive atom so that Mw of peptide is not affected and the γ radiation is then measured as a function of tissue depth. Computer software is employed to create a three-dimensional image of the distribution of the substance in brain and other tissues **[37]**.

An important outcome of this study is that the selected pool of bromelain peptides not only irreversibly dissociates preformed Aβ fibrils (Fig. 3), but also inhibits formation of Aβ fibrils from monomeric and oligomeric states. This can be achieved in two ways; first, bromelain peptides may bind with monomeric Aβ peptide presumably protecting the hydrophobic site/s of interaction leading to prevention of aggregation or second, upon binding with bromelain peptides, Aβ peptide undergoes an irreversible conformation change whereby potency of self-association is lost. It is now well established that the following equilibration exists during the process or even after fibril formation **[38]**:

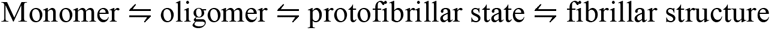

There are evidences from immunological studies using antibodies specific to monomeric or oligomeric Aβ peptides that even after fibril formation, monomers and oligomers do exist in equilibrium **[39].** Certainly the equilibrium shifts to the right (towards fibril formation) when stable fibrillar structures are formed while remains in the middle during onset of aggregation. Fibrillar structures being very stable, it is difficult to conceive that bromelain peptides directly interact with them and make them unstable. On the other hand, bromelain peptides irreversibly inhibit formation of fibrillar structures from monomeric Aβ peptide indicating positive interaction leading to conformation change. This is demonstrated indirectly from interaction with the fluorophore ANS (Fig. 4A-4B) and directly from CD and FT-IR spectra (Fig. 4C–4F).

An intriguing part of these studies is that often requirement of inhibitor peptides is sub-stoichiometric with respect to Aβ peptides. Though it is not possible to predict molecularity of an aggregate or a mixture of oligomers or peptides in a pool, concentrations of Aβ monomer and synthetic peptides could be accurately determined. From this information, it can be predicted that formation of a stable [Aβ monomer–synthetic peptide] binary complex of 1:1 stoichiometry leading to inhibition of aggregate formation is not possible. The event most pertinent to this situation is synthetic or bromelain derived peptides bind with Aβ monomer causing irreversible conformation change. As a result, after dissociation of the binary complex, Aβ peptides permanently lose their ability to form oligomers and fibrillar structures. The inhibitor peptide in its free state after dissociation from the complex recycles the reaction. This is similar to enzyme turnover.

A large number of medicinal properties have been attributed to fruit ‘bromelain’–the fresh fruit extract of pineapple. Though cysteine protease bromelain is the major constituent of the extract, other accompanying components are peroxidases, acid phosphatases, glycosidases, ribonucleases, cellulases, glycoproteins, carbohydrates, protease inhibitors together with organic and inorganic compounds. It is believed that a wide array of medicinal properties reside in this battery of enzymes **[14]**. Pineapple is a seasonal crop. The fruit is consumed more as a processed product rather than in its raw form. We have reported earlier that due to harsh sterilization conditions, these products are completely devoid of enzymatic activities as the enzymes are either thermally denatured or proteolytically degraded **[11]**. Since peptides formed after extensive degradation of ‘bromelain’ enzymes can destabilize fibrillar structures as reported in this study, processed pineapple should be capable of performing similar functions. The statement of Greek physician and philosopher, Hippocrates, ‘let food be thy medicine and medicine be thy food’, appears meaningful.

## CONCLUSION

Peptides generated from fruit bromelain under human digestive conditions can inhibit aggregate formation from monomeric state of amyloidogenic peptides Aβ40/42 besides facilitating destabilisation of preformed amyloid fibrils *in vitro. Ex vivo* studies using PC12 neuronal cells and *in vivo* studies using animal models suggested reversal of neurotoxicity caused by amyloid peptides in presence of bromelain derived peptides. Probable underlying mechanism has been proposed. Since pineapple is edible, one can initiate clinical trials to determine preventive effects of neuronal toxicities in AD patients, elderly individuals and in subjects with mild cognitive impairment using fruit peptides.

## ACKNOWLEDGEMENTS

This research was partially supported by CSIR Network Project (mIND BSC 0115). Das S. was supported from the same source. Dutta S was funded by UGC-SRF and RKP was supported by CSIR. We thank Dr. Krishnananda Chattopadhyay and Dr. Sandhya Rekha Dungdung of CSIR-IICB for providing their DLS and cell culture facility respectively. Special thanks to Mr. T.Muruganandan, Mr. Sailen De, Mr. Jishu Mondal, Mr. Satyabrata Samaddar, Mr, Sandip Chakraborty and Mr. Diptendu Bhattacharya are due for their technical support in AFM, TEM, CD, FT-IR, MALDI and ESI-MS facilities respectively.

## AUTHOR CONTRIBUTIONS

Conceptualization, S.Das, S.Dutta, R.K.P, S.C.B, U.C.H. and D.B.; Investigation, S.Das, S.Dutta. and R.K.P; Data Analysis, S.Das, S.Dutta. and R.K.P,; Writing, S.Das., S. Dutta. and D.B.; Funding Acquisition, Resources, and Supervision, S.C.B., U.C.H and D.B.

## DATA SHARING AND DATA ACCESSIBILITY

The data that support the findings of this study are available from the corresponding author upon reasonable request.

## SUPPLEMENTAL INFORMATION

### Supplemental Figures

**Fig S1:**
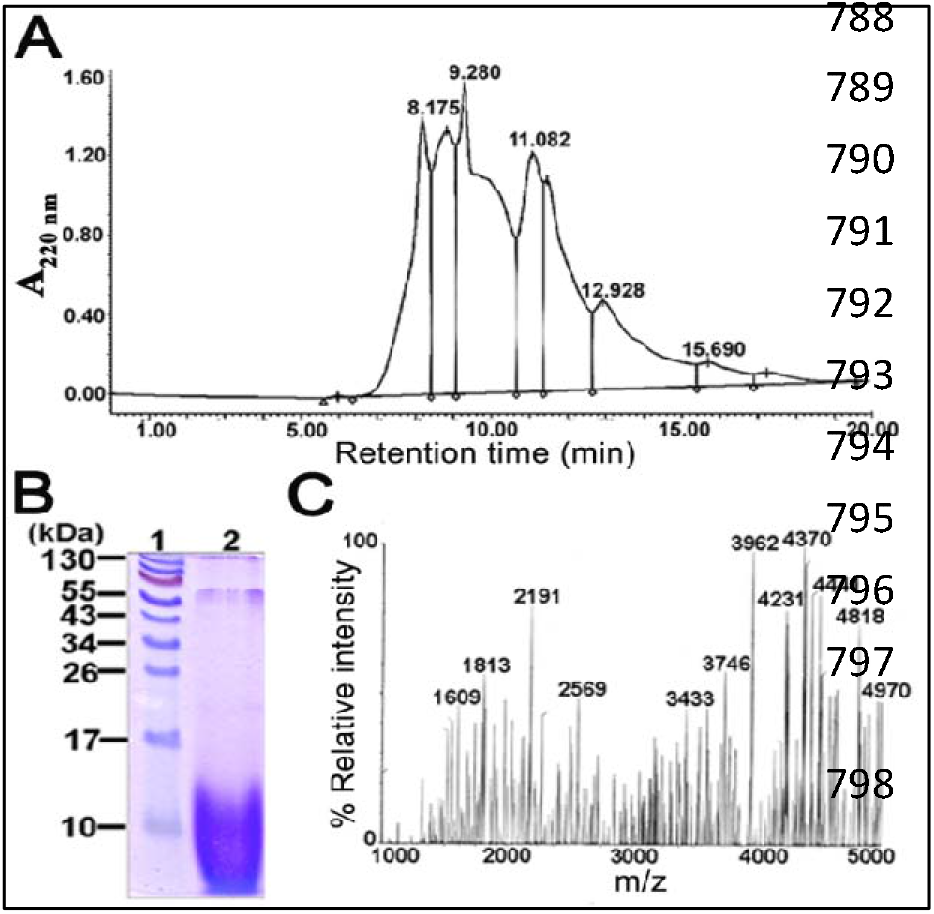
Purification and mass analysis of bromelain peptide pool. (A) SE-HPLC of fruit bromelain derived peptide pool using a Waters Protein Pak 60 column equilibrated with 10 mM Na-P buffer (pH 7.5). Flow rate was 0.8 ml/min and elution was followed at 220 nm. Retention times of major peaks have been indicated. (B) 15 % SDS-PAGE profiles. Lane 1, Protein markers; Lane 2, Peptide pool marked by bar in the chromatogram. (C) MALDI-MS spectrum of SE-HPLC eluted pool of the peptides. Related to Fig. 2.

**Fig S2:**
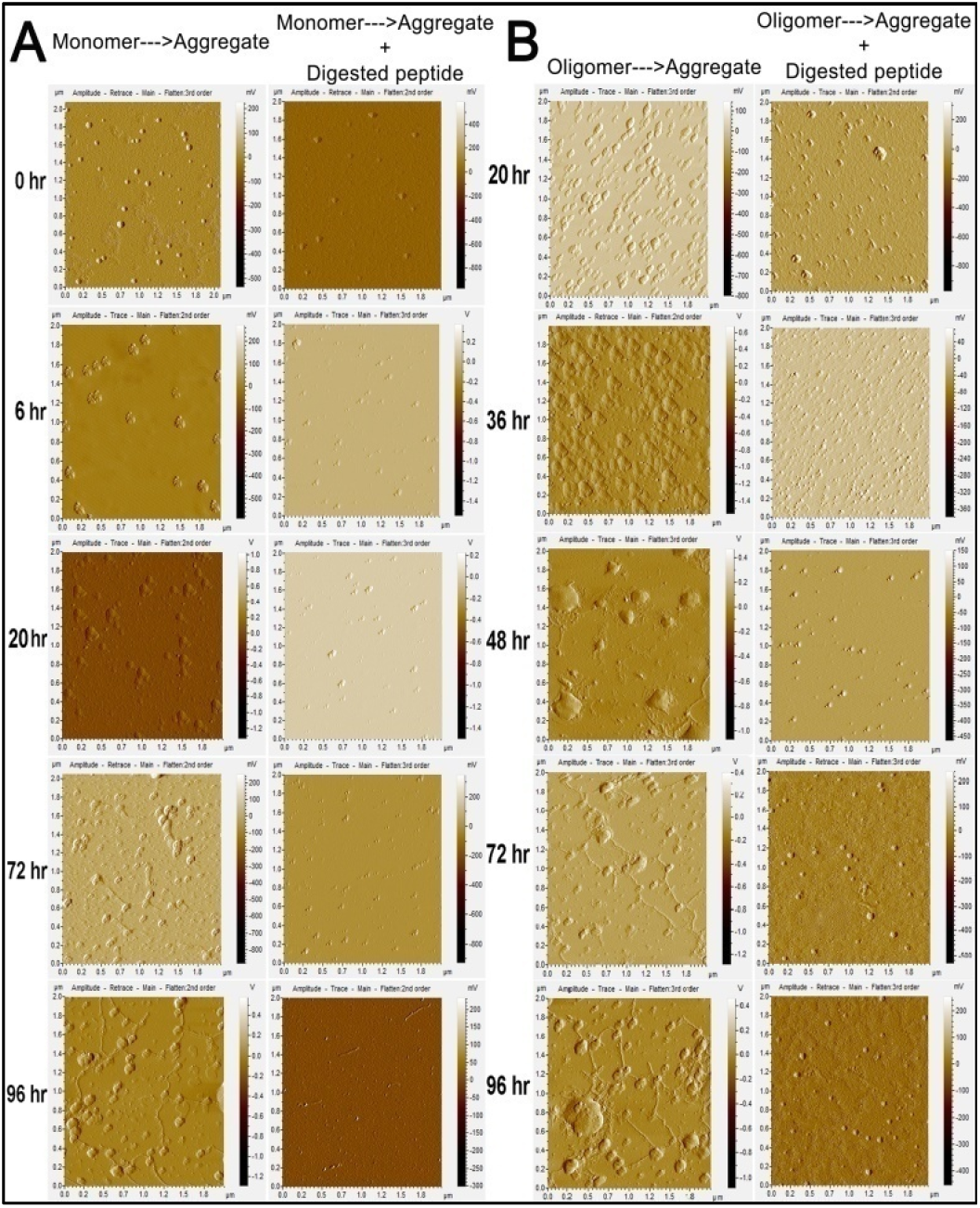
Time course of aggregation as followed by AFM micrographs. (A) (Monomers to aggregates): Aβ (1 – 40) co-incubated in the absence or presence of 7 μM of digested peptide at 37 °C in Na-phosphate buffer (pH 7.5). The aliquots were taken at 0, 12, 24, 72 and 96 hr and analyzed for the presence or absence of aggregates (B) oligomers to aggregates: Aβ 40 was incubated for 20 hr under fibrillating conditions and digested peptide was added after 20 hrs of incubation. Aliquots were taken at 20, 36, 48, 72 and 96 hr for analysis of the presence or absence of aggregate. Related to Fig. 3.

